# The Mn-motif protein MAP6d1 assembles ciliary doublet microtubules

**DOI:** 10.1101/2025.02.17.638590

**Authors:** Dharshini Gopal, Juliette Wu, Julie Delaroche, Christophe Bosc, Manon De Andrade, Eric Denarier, Gregory Effantin, Annie Andrieux, Sylvie Gory-Fauré, Laurence Serre, Isabelle Arnal

## Abstract

Most eukaryotic cells have cilia that serve vital functions in sensing, signaling, motility. The core architecture of cilia is an array of microtubule doublets, which consist of a complete A-tubule and an incomplete B-tubule. The mechanisms governing the assembly of this complex structure remain poorly understood. Here, using total internal reflection fluorescence microscopy and cryo-electron tomography, we investigate the role of MAP6d1, a brain-specific protein containing microtubule lumen-targeting Mn-motifs. We show that MAP6d1 assembles stable microtubule doublets by recruiting tubulin dimers onto the lattice of the A-tubule to initiate the nucleation of the B-tubule. MAP6d1 also promotes the formation of luminal protofilaments in singlet and doublet microtubules, a previously undescribed phenomenon that likely enhances microtubule stability. In neurons, MAP6d1 localises to the proximal part of primary cilia via its Mn-motif, with its loss resulting in shortened cilia, a characteristic of ciliopathies. MAP6d1 is thus the first microtubule-associated protein found to assemble microtubule doublets, uncovering new functions for Mn-motif proteins in neurons.

## Introduction

Microtubules are one of the major classes of cytoskeletal proteins that serve a wide range of cellular functions, providing tracks for cargo transport and structural support to maintain cellular architecture and enable motility^1,2^. In order for cells to quickly reorganise their cytoskeleton in response to various triggers, dynamic microtubules alternate between phases of growth and shrinkage, known as dynamic instability^3^. In contrast, long-lived stable microtubules with reduced tubulin turnover predominate in terminally differentiated cells such as neurons, where they maintain cell morphology and support long-distance intracellular transport^4^. Particularly stable microtubules are also found in the axonemes of cilia and flagella, where they form complex arrays of doublets, ensuring the integrity and functions of these cellular appendages^5^. Motile cilia and flagella are responsible for moving cells (e.g., sperm) or a part of their environment (e.g., mucus along the trachea), while non-motile or primary cilia function as antenna for sensory perception and signal transduction on most vertebrate cells^6,7^. Defects in cilia assembly and structure can result in a number of diseases, collectively referred to as ciliopathies^8,9^.

Cilia share a core structure, the axoneme, composed of nine doublet microtubules arranged radially around two central singlet microtubules in motile cilia, or without a central pair in primary cilia. Each doublet microtubule typically comprises one complete 13 protofilament A-tubule and one incomplete 10 protofilament B-tubule. Studies of cilia and flagella in unicellular organisms such as *Chlamydomonas* and *Tetrahymena* revealed the presence of proteins located in the microtubule lumen known as Microtubule Inner Proteins (MIPs), which have since emerged as key contributors to the remarkable stability of axonemal doublet microtubules^10–14^. In recent years, more than 60 axonemal MIPs have been identified across species, and they exhibit a variety of luminal microtubule-binding modes^15–17^. To date, however, the function of most MIPs remains obscure. More generally, the mechanisms by which microtubule doublets are assembled and the role of MIPs in this process are still poorly understood.

MIPs are also present in neurons, where they have long been observed as intraluminal particles in both cytoplasmic and ciliary microtubules^18–23^. By analogy with axonemal MIPs, they are thought to contribute to neuronal microtubule stability, although their identity remains largely unknown. MAP6, a microtubule-stabilising factor linked to psychiatric disorders^24,25^, was the first neuronal MIP located in the microtubule lumen, where it generates highly stable microtubules that grow in a helicoidal pattern^26^. MAP6 belongs to the SAXO (Stabiliser of AXOnemal microtubules) family of proteins that contain a small helical motif in their microtubule-binding domains known as the Mn-motif^17,27–29^. This motif has recently been recognised as a universal microtubule luminal binding motif in axonemal MIPs, highlighting its potential significance in stabilising both axonemal and neuronal microtubules. Interestingly, only two SAXO family members, MAP6 and MAP6d1, are currently known to be expressed in the brain.

Here we focus on MAP6d1, the sole SAXO family member known to be expressed exclusively in the postnatal brain^30^. MAP6d1 contains two Mn-motifs and can protect microtubules from drug- and cold-induced depolymerisation via its Mn-motif-containing microtubule-binding domains^31^. Beyond its cellular localisation to microtubules, mitochondria, Golgi and membranes following palmitoylation^30^, little is known about MAP6d1 functions. To investigate the effects of MAP6d1 on microtubules and related cellular roles, we used a combination of Total Internal Reflection Fluorescence (TIRF) microscopy-based *in vitro* reconstitution assays, cryo-electron tomography (cryo-ET), and cultured hippocampal neurons. Our findings uncover a direct role for MAP6d1 in the assembly and stabilisation of doublet microtubules, shedding light on its involvement in maintaining the architecture of neuronal primary cilia.

## Results

### MAP6d1 stabilises microtubules by inducing pauses

To gain insight into the mechanism underlying MAP6d1’s ability to stabilise microtubules, we used TIRF microscopy and *in vitro* reconstitution assays to study microtubule dynamics (**Fig. 1a** and **Supplementary Movie 1**). We found that MAP6d1 did not affect the frequency of catastrophes but did reduce both the growth and shrinkage rates while promoting rescues, at both plus and minus ends (**Fig. 1b** and Supplementary Fig. 2a). The net effect was the occurrence of pauses during which the microtubules neither grew nor shrank. Pauses became more frequent at MAP6d1 concentrations of 20 nM and lasted significantly longer at concentrations of 50 nM (**Fig. 1c**). The longest pauses, lasting up to 20 minutes, blocked microtubule dynamics, with a mean paused microtubule length of about 8 ± 4.2 *μ*m (n = 45 microtubules).

**Figure 1:**
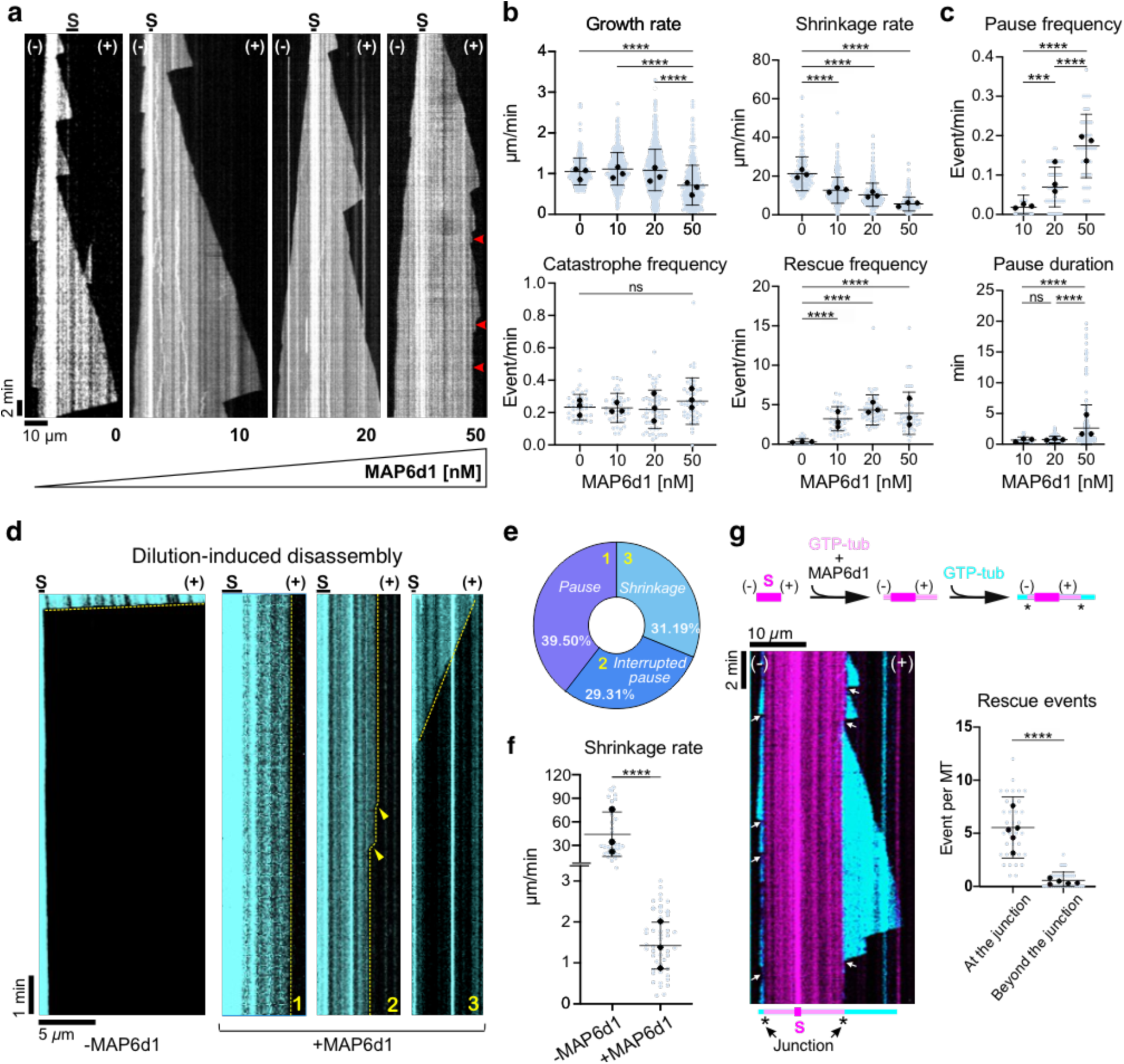
MAP6d1 stabilises microtubules by inducing pauses. **a** Representative kymographs of microtubules grown from seeds (S) and 12 *μ*M tubulin with increasing MAP6d1 concentrations. Arrowheads indicate pauses. **b** Graphs showing the growth rate (n = 301, 427, 712, and 364 events for 0, 10, 20 and 50 nM MAP6d1, respectively), shrinkage rate (n = 168, 274, 334, and 264 events), catastrophe and rescue frequencies (n = 36, 41, 54 and 45 microtubules) of microtubules assembled with or without MAP6d1. **c** Graphs showing the pause frequency (n = 41, 54 and 45 microtubules, for 10, 20 and 50 nM MAP6d1 respectively) and durations (n = 24, 121 and 245 events) in similar conditions. In the absence of MAP6d1, microtubules did not exhibit any pauses. **d** Representative kymographs of microtubules undergoing disassembly after tubulin dilution. Microtubules were grown from seeds (S) and 25 *μ*M tubulin (control, *left*) or 12 *μ*M tubulin and 50 nM MAP6d1 (*right*). After dilution, three populations were observed in the presence of MAP6d1: continuous pause after dilution, pauses interrupted by small catastrophes (arrowheads), and continuous slow depolymerisation. **e** Percentages of the three populations presented in **(D). f** Shrinkage rate of control microtubules and microtubules of population 3 shown in **D** (n = 32 and 50 events, respectively). **g** Kymograph showing rescue events of a microtubule grown with 12 µM GTP-tubulin (cyan) at the extremity of a microtubule assembled with 12 µM GTP-tubulin and 50 nM MAP6d1 (magenta). The experimental procedure is depicted above the kymograph. The kymograph shows the behaviour of a dynamic microtubule recorded after perfusion of GTP-tubulin. Arrows indicate rescue events occurring at the junction (*) between the MAP6d1-assembled microtubule (magenta) and the newly growing dynamic microtubule (cyan). The graph represents the number of rescue events occurring at or beyond the junction (n = 35 microtubules). For graphs in B, C, F and G: bars represent mean ± SD from at least three independent experiments. Circles with different colours represent the mean of each individual experiment. ***p<0.001, ****p<0.0001, ns = non-significant, Kruskal-Wallis analysis of variance (ANOVA) followed by post hoc Dunn’s multiple comparisons tests.

To evaluate the stability of these MAP6d1-paused microtubules, we performed dilution-induced depolymerisation assays. Microtubules assembled with tubulin alone quickly depolymerised within seconds upon buffer perfusion (**Fig. 1d**, left), whereas those assembled in the presence of MAP6d1 resisted depolymerisation with three distinct behaviours (**Fig. 1d**, right). One population of microtubules (39.5%) remained in a paused state for the entire duration of the movie (30 min) following buffer perfusion; a second (29.3%) showed pauses interrupted by small catastrophes; and a third population (31.2%) continuously but slowly depolymerised after buffer perfusion (**Fig. 1e**). The shrinkage rate of these slowly depolymerising microtubules was 50-fold slower than that observed for microtubules assembled without MAP6d1 (**Fig. 1f**).

We next performed a two-step perfusion TIRF experiment, polymerising microtubules from red tubulin in the presence of MAP6d1 then exchanging the solution with green-fluorescent tubulin before recording. In this experiment, approximately 70% of the dynamic green-fluorescent microtubules, polymerising from the extremities of MAP6d1 co-assembled microtubules, depolymerised up to the junction with the MAP6d1-assembled microtubule lattice, which served as a rescue point (**Fig. 1g**).

Thus, microtubules assembled in the presence of MAP6d1 resist buffer-induced depolymerisation and are stable enough to stop the depolymerisation of a dynamic microtubule.

Overall, these results reveal that MAP6d1 is a strong microtubule stabiliser.

### MAP6d1 induces the assembly of doublet microtubules by recruiting tubulin

To investigate how MAP6d1 influences microtubule architecture, we performed negative staining electron microscopy on microtubules polymerised in the presence or absence of MAP6d1 (**Fig. 2a**). To our surprise, MAP6d1 enabled the formation of doublet microtubules, which were not seen with microtubules polymerised with tubulin alone. Cryo-ET revealed distinct microtubule architectures, including doublet microtubules with varying B-tubules, as shown in tomograms in (**Fig. 2b** and **Supplementary Fig. 3**). The complete microtubules, i.e., the A-tubules of doublet microtubules in our dataset, were consistently composed of 14 protofilaments, whereas the z-slices of the example tomograms delineated 11 classes of B-tubule with different numbers of protofilaments. Most B-tubules had 7 to 11 protofilaments, with 10 protofilaments being the most abundant class (∼40 % of the extracted particles). This heterogeneity in B-tubule architecture could represent different stages of doublet microtubule assembly. Notably, the B-tubule was highly flexible, as its extremity did not anchor back to the A-tubule but exhibited global curvature similar to that observed *in vivo* (**Fig. 2c**). Using subtomogram averaging, we reconstructed a doublet microtubule from these data showing an A-tubule with 14 protofilaments and a B-tubule with 10 protofilaments (**Fig. 2d**; see methods).

**Figure 2:**
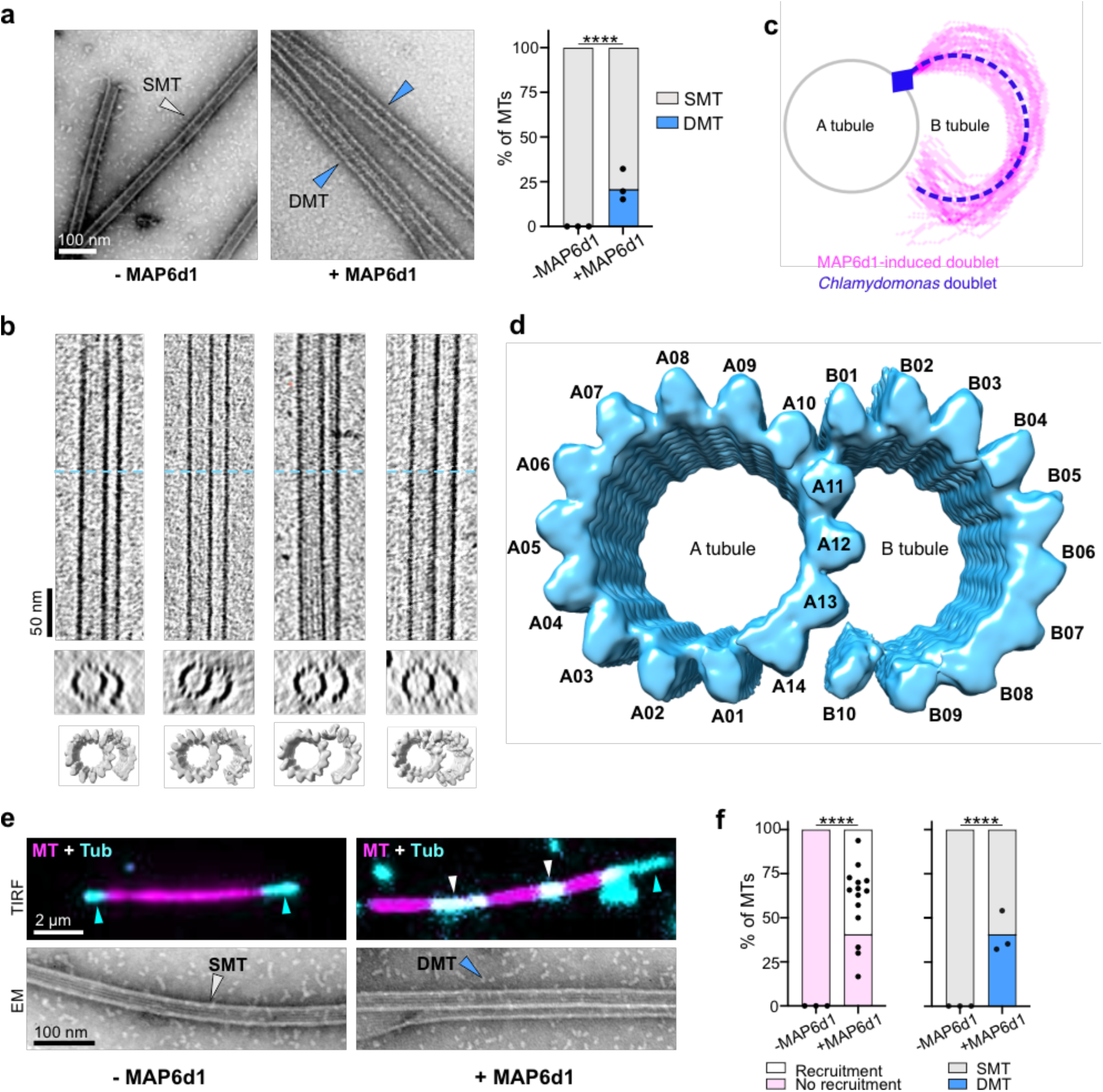
MAP6d1 induces the assembly of doublet microtubules by recruiting tubulin. **a** Negative staining electron microscopy images of microtubules assembled with 25 *μ*M tubulin in the absence or presence of 250 nM MAP6d1. White and blue arrowheads indicate singlet microtubules (SMTs) and doublet microtubules (DMTs) respectively. The graph represents the percentage of SMTs and DMTs assembled without and with MAP6d1 (total measured lengths of 304 and 401 µm, respectively). **b** Examples of cryo-electron tomograms of doublet microtubules assembled with MAP6d1. The top panel shows longitudinal sections of tomograms along with a transverse section at the position of the blue dashed lines. A subtomogram average map is presented below each tomogram. **c** Traces of the B-tubule of MAP6d1-induced doublet microtubules starting at the outer junction (blue square) and overlapping the curvature of the B-tubule of the *Chlamydomonas* flagellar doublet microtubule (EMD:40621) for comparison (n = 45 B-tubule curvatures). **d** Reconstruction of MAP6d1-induced doublet microtubules obtained by combining subtomogram averaging models obtained by individually masking the A- and the B-tubules. **e** *Top*: snapshots of TIRF microscopy recorded movies of 0.325 *μ*M tubulin (cyan) mixed with GMPCPP-stabilised microtubule seeds (magenta) in the absence or presence of 100 nM MAP6d1 and 50 *μ*M GMPCPP. White and blue arrowheads indicate tubulin recruitment and polymerisation at microtubule ends, respectively. *Bottom*: negative staining electron microscopy images of microtubules assembled in the same conditions as for TIRF microscopy. White and blue arrowheads indicate SMT and DMT, respectively. **f** *Left*: the percentage of microtubules that have or have not recruited tubulin. The total number of analysed microtubules is 57 and 282 for the control and with MAP6d1, respectively (from at least three independent experiments). Black dots represent the percentage of microtubules with recruitment for each experiment. *Right*: percentage of singlet and doublet microtubules in the control and with MAP6d1. The total measured lengths of microtubules were 225 and 543 *μ*m for the control and with MAP6d1, respectively (three independent experiments). Black dots represent the percentage of doublet microtubules for each experiment. For graphs in A and G: ****p<0.0001, Fischer’s exact contingency test.

To better understand the underlying mechanism of doublet microtubule formation, we performed tubulin recruitment assays using TIRF microscopy (**Fig. 2e**). Red-fluorescent pre-polymerised microtubules were incubated with free green-fluorescent tubulin. In the absence of MAP6d1, there was no tubulin recruitment (**Fig. 2e**, *left*), but with MAP6d1 we observed recruitment of free tubulin onto 57% of the pre-polymerised microtubules (**Fig. 2e, f**). We also noticed elongation of this recruited tubulin along the pre-existing microtubules (**Supplementary Movie 2**). To investigate the impact of tubulin recruitment on microtubule structure, we next imaged these samples by negative staining electron microscopy. In the presence of MAP6d1, 40% of microtubules were doublets, but without MAP6d1, there appeared only singlet microtubules (**Fig. 2e, f**). This confirms that the ability of MAP6d1 to assemble doublet microtubules depends on its capacity to recruit tubulin dimers along the microtubule lattice.

### The Mn-motif of MAP6d1 determines its microtubule-regulating activities

We analysed the domains previously identified as essential for properties of MAP6d1 **(Supplementary Fig. 1**). The mutant MAP6d1-1′Mn2, which deletes the microtubule-binding domain encompassing the second Mn-motif and its C-terminal flanking region, conserved in MAP6^30^, abolished the microtubule-stabilising activity of MAP6d1 (**Fig. 3a, b** and **Supplementary Fig. 2b**). This deletion also eliminated the ability of MAP6d1 to recruit tubulin dimers along the microtubule lattice and to promote microtubule doublet assembly (**Fig. 3c, d**). To assess whether these effects were directly linked to the Mn-motif, we designed a mutant of MAP6d1 in which the seven residues of the second Mn-motif were replaced by alanine (MAP6d1-Mn2-7A). This mutant was also unable to stabilise microtubules, promote doublet assembly or recruit tubulin onto the microtubule lattice (**Fig. 3**). The second Mn-motif clearly contributes to the observed microtubule-regulatory roles of MAP6d1.

**Figure 3:**
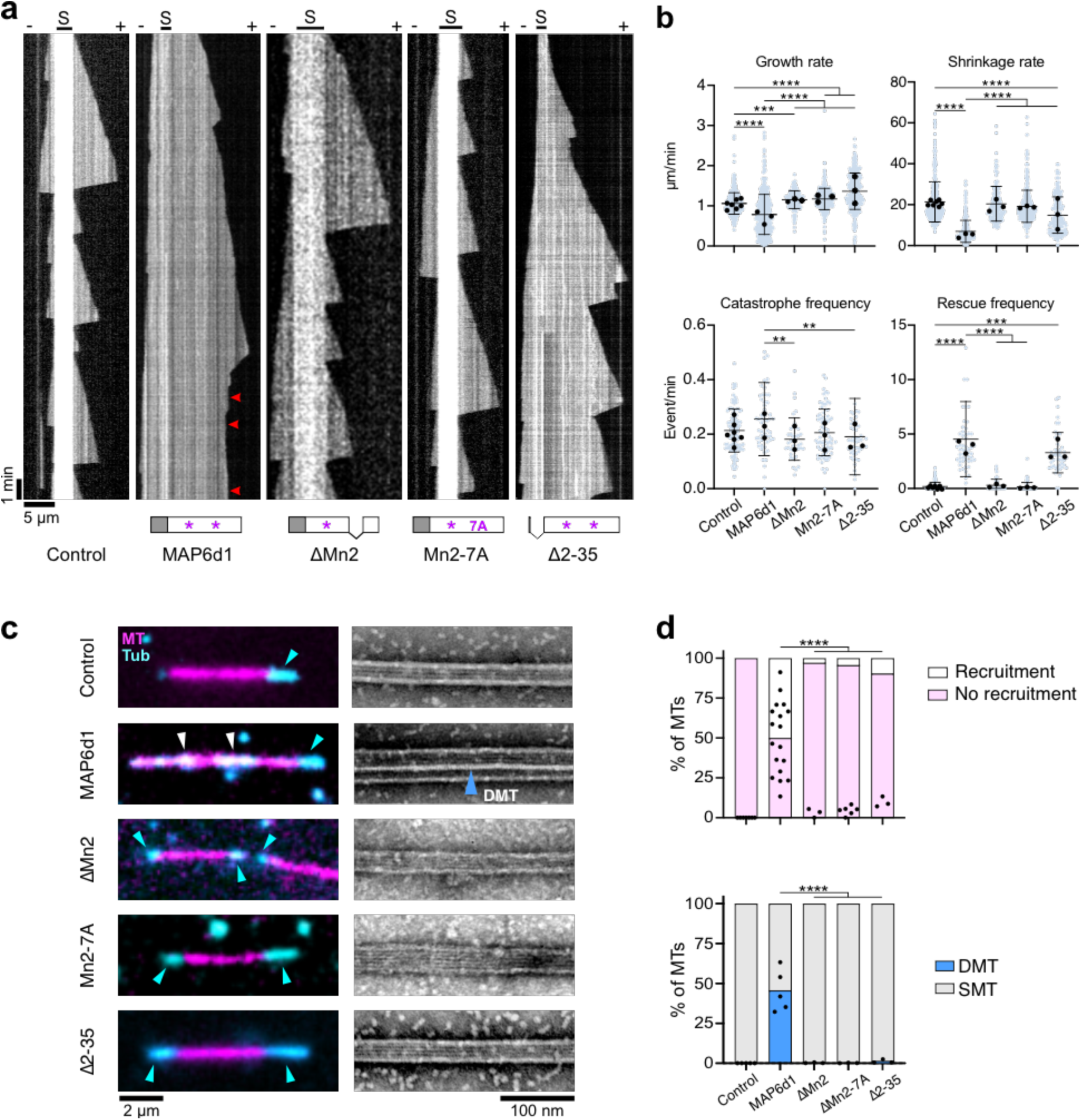
The microtubule-regulating activities of MAP6d1 depend on its Mn-motif and N- terminus. **a** Kymographs depicting microtubules grown from seeds (S) and 12 *μ*M tubulin alone or with 50 nM MAP6d1, 300 nM MAP6d1-ΔMn2, 300 nM MAP6d1-Mn2-7A or 50 nM MAP6d1-Δ2-35 (n = 105, 57, 81, 49 and 38 microtubules, respectively). MAP6d1-ΔMn2/-Mn2-7A concentrations were increased to 300 nM because of their lower microtubule-binding activity relative to MAP6d1 (**Supplementary Fig. 1b, c**). Arrowheads indicate microtubule pauses. **b** Dynamical parameters of microtubules assembled with tubulin alone and in the presence of MAP6d1, MAP6d1-ΔMn2 MAP6d1-Mn2-7A and MAP6d1-Δ2-35: growth rate (n = 556, 437, 190, 397 and 286 events, respectively), shrinkage rate (n = 444, 284, 162, 348 and 197 events, respectively), catastrophe and rescue frequencies (n = 105, 56, 38, 81 and 48 microtubules, respectively). Bars represent mean ± SD from at least three independent experiments. Circles with different colours represent the mean of each individual experiment. **p<0.01, *** p<0.001, **** p<0.0001. Kruskal-Wallis analysis of variance (ANOVA) followed by post hoc Dunn’s multiple comparisons tests (only significant statistics are indicated). **c** *Left*: snapshots of TIRF movies of 0.325 *μ*M tubulin (cyan) mixed with GMPCPP seeds (magenta) in the absence or presence of 100 nM MAP6d1, 300 nM MAP6d1-ΔMn2 300 nM MAP6d1-Mn2-7A and 100 nM MAP6d1-Δ2-35. White and blue arrowheads indicate tubulin recruitment and polymerisation at the seed extremities, respectively. *Right*: Negative staining electron microscopy images of microtubules assembled in the same conditions as for TIRF microscopy. The blue arrowhead indicates a doublet microtubule. **d** *Top*: percentage of microtubules with or without recruited tubulin (57, 282, 63, 164 and 152 microtubules for the control and with MAP6d1, MAP6d1-ΔMn2, MAP6d1-Mn2-7A and MAP6d1-Δ2-35, respectively). *Bottom*: percentage of singlet and doublet microtubules in the control and with MAP6d1 and its mutants (total measured lengths of 361, 752, 149, 221 and 198 *μ*m for MAP6d1, MAP6d1-ΔMn2, MAP6d1-Mn2-7A and MAP6d1-Δ2-35, respectively). Black dots represent the percentage of doublet microtubules for each experiment. ****p<0.0001, Fischer’s exact contingency test. All statistical analyses were performed from at least three independent experiments.

### The N-terminal domain of MAP6d1 is required for microtubule pausing and doublet formation

Next, we compared the microtubule-regulatory activity of MAP6d1 to that of a mutant lacking its N-terminal domain (MAP6d1-Δ2-35), which is conserved in MAP6 and involved in MAP6-mediated microtubule stabilisation and luminal particle formation^26,30^. MAP6d1-Δ2-35 still stabilised microtubules by increasing the rescue frequency and reducing both the shrinkage rate and catastrophe frequency (**Fig. 3a, b** and **Supplementary Fig. 2b**), leading to persistent microtubule growth. Unlike MAP6d1, however, MAP6d1-Δ2-35 did not induce microtubule pausing (**Fig. 3a**) and it suppressed doublet microtubule formation (only ∼1% vs. the ∼40 % with MAP6d1; **Fig. 3c, d**). Consistent with these observations, this mutant weakly recruited free tubulin along stabilised microtubules (∼9 % versus ∼57 % for MAP6d1) (**Fig. 3c, d**). Thus, although the N-terminal domain is not needed for stabilising microtubules, it is essential to induce pauses and to form doublets by recruiting tubulin.

### MAP6d1 localises to microtubule doublets in neuronal primary cilia and regulates ciliary length

Building on evidence of MAP6d1’s ability to assemble microtubule doublets *in vitro*, we moved to studies in neurons, focusing on primary cilia. We compared primary cilia from cultured hippocampal neurons derived from Wild-Type (WT) and MAP6d1-KnockOut (KO) mice (**Fig. 4a, b**). The percentage of ciliated neurons were similar in both cases (**Supplementary Fig. 4a**). At 4 days *in vitro* (DIV), ciliary length (4.3 ± 1.861 *μ*m) was also similar to that observed in WT (4.4 ± 1.746 *μ*m) (**Fig. 4a, b**). Given that MAP6d1 expression begins around 7 DIV^30^, we re-examined ciliary length at 9 DIV, and found that MAP6d1-KO cilia had become 15% shorter than those in WT neurons (5.38 ± 2.022 *μ*m to 4.7 ± 1.695 *μ*m); this difference persisted through 23 DIV (**Fig. 4b** and **Supplementary Fig. 4b**).

**Figure 4.**
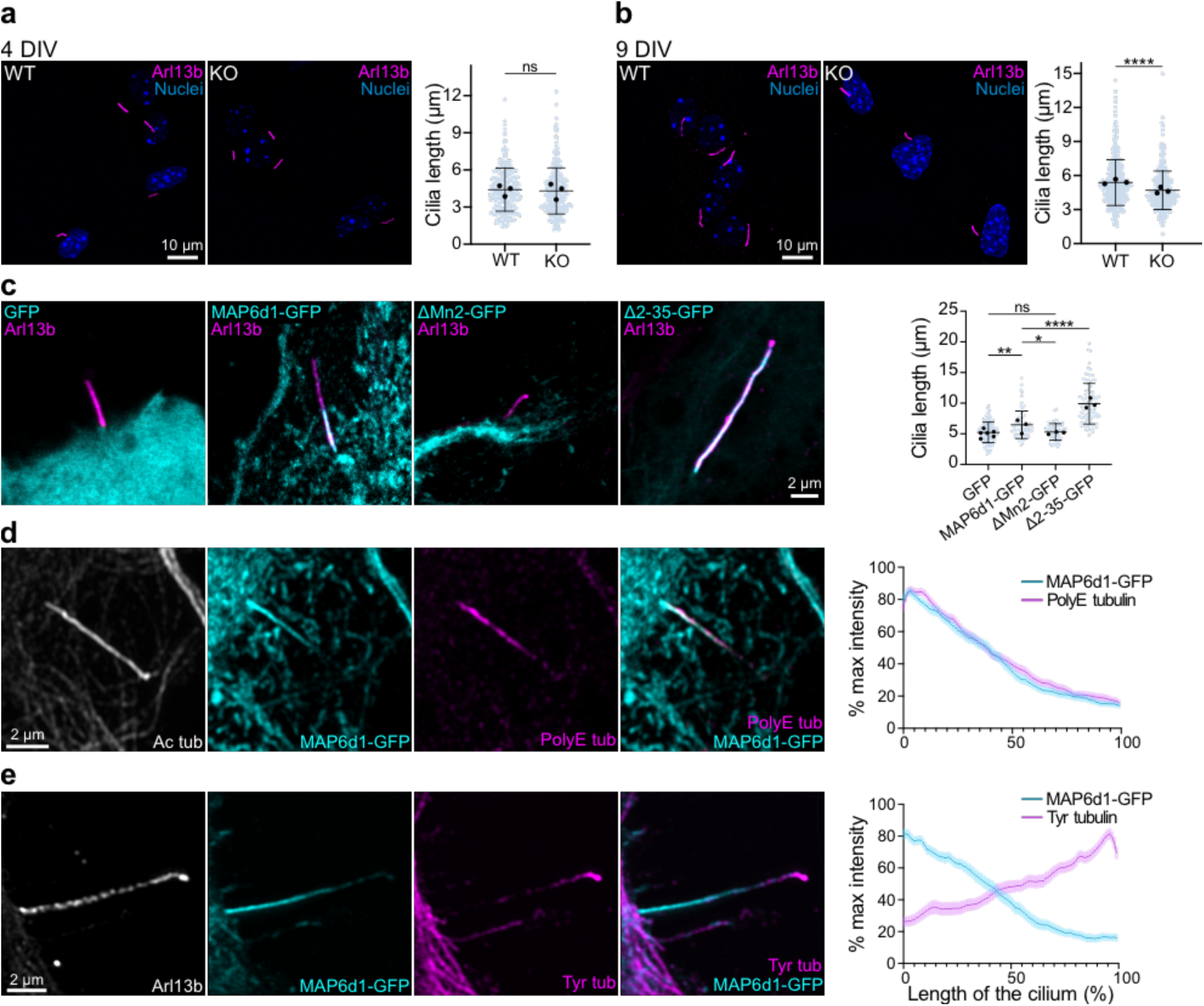
MAP6d1 preferentially localises to microtubule doublets in neuronal primary cilia and regulates ciliary length. **a** Representative images of 4 DIV WT and MAP6d1-KO hippocampal neurons stained against cilia marker Arl13b and nuclear dye Hoechst. The graph represents the cilia length of WT (n = 253) and KO neurons (n = 262). **b** Representative images of 9 DIV WT and MAP6d1-KO hippocampal neurons stained against cilia marker Arl13b and nuclear dye Hoechst. The graph represents the cilia length of WT (n = 407) and KO neurons (n = 398). For graphs in (**a**) and (**b**), bars represent mean ± SD from three independent experiments. Circles with different colours represent the mean of each individual experiment. ns = not significant, ****p < 0.0001, by Mann-Whitney’s test. **c** The localisation of ectopic GFP, MAP6d1-GFP, MAP6d1-ΔMn2-GFP and MAP6d1-Δ2-35-GFP in comparison to ciliary marker Arl13b, in 7 DIV hippocampal neurons. The graph represents the cilia length in neurons transfected with plasmids encoding GFP (n = 128), MAP6d1-GFP (n = 68), MAP6d1-ΔMn2-GFP (n = 59), MAP6d1-Δ2-35-GFP (n = 86) from at least three independent experiments. Graph presents individual data points with bars representing mean ± SD. Circles with different colours represent the mean of each individual experiment. ns = not significant, *p<0.05, ** p<0.001, ****p<0.0001 by by one-way ANOVA and pairwise comparisons. **d** Representative images of primary cilia from 7 DIV hippocampal neurons expressing ectopic MAP6d1-GFP and stained against acetylated (Ac tub) and polyglutamylated tubulin (PolyE tub). The graph represents the percentage of normalised maximum fluorescence of MAP6d1-GFP and polyglutamylated tubulin (PolyE tub) along the ciliary length determined by acetylated tubulin (Ac tub). The values represent mean ± SEM of n = 58 cilia from 3 independent experiments. **e** Representative images of primary cilia from 7 DIV hippocampal neurons expressing ectopic MAP6d1-GFP stained against Arl13b and tyrosinated tubulin (Tyr tub). The graph represents the percentage of normalised maximum fluorescence of MAP6d1-GFP and tyrosinated tubulin along the ciliary length determined by Arl13b. The bars represent mean ± SEM of n = 46 cilia from 3 experiments.

To determine the localisation of MAP6d1 in neurons, we expressed ectopic GFP-tagged MAP6d1 and the ΔMn2 and Δ2-35 mutants in the hippocampal cells. MAP6d1-GFP localised to the proximal part of the primary cilia in all transfected cells and led to cilia ∼24% longer than those in control cells (**Fig. 4c** and **Supplementary Fig. 4c**). This asymmetric distribution contrasted with that of the ciliary marker Arl13b, which spans the entire length of the cilium. MAP6d1-ΔMn2-GFP failed to localise to the cilia, which were similar in length to those in wild-type cells. On the other hand, MAP6d1-Δ2-35-GFP was present all along the cilia, which were twice as long as those in control neurons (**Fig. 4c**). MAP6d1’s ciliary localisation therefore requires the Mn2 motif, and its specific proximal ciliary targeting is linked to the N-terminal domain.

The proximal localisation of MAP6d1 corresponds to the microtubule doublet region identified by cryo-ET showing that doublet microtubules lose their B-tubule, eventually leaving only singlet A-tubules in the distal part of the cilia^32,33^. To further investigate MAP6d1’s preferential proximal enrichment, we performed co-immunofluorescence staining with post-translational modifications (PTMs) of tubulin known to accumulate in ciliary axonemes. We found that MAP6d1-GFP co-localised with polyglutamylated tubulin (**Fig. 4d**), a PTM enriched on the B-tubules in motile cilia^34–36^. Conversely, MAP6d1-GFP and tyrosinated tubulin, a PTM associated with dynamic microtubules^37,38^, display opposite localisation patterns, with tyrosinated tubulin being enriched at the distal end of the cilia (**Fig. 4e**). This finding is consistent with previous observations that single dynamic microtubules are found predominantly at the ciliary tips^39–41^.

These data indicate that MAP6d1 plays a role in assembling and stabilising microtubule doublet architecture in neuronal cilia.

### MAP6d1 assembles protofilaments in the lumen of singlet and doublet microtubules

Tomograms of microtubules assembled in the presence of MAP6d1 revealed peculiar structures within the lumen of both doublet and singlet microtubules (**Fig. 5a**). These structures correspond to two adjacent tubulin protofilaments that would interact with the microtubule internal surface. Indeed, subtomogram averages applied to these MAP6d1-induced structural entities allowed to obtain reconstructions of doublet and singlet microtubules with two luminal protofilaments (**Fig. 5b, c**). These reconstructions indicate that the luminal protofilaments in the A-tubule of the doublet are located facing protofilaments A10 to A13, adjacent to the putative seam (**Fig. 5c**). We did not detect any obvious lattice defects in proximity to these luminal protofilaments (in a total of 220 microtubules), suggesting that luminal protofilaments are formed during microtubule copolymerisation with MAP6d1 rather than through diffusion via breaks in the lattice.

**Figure 5.**
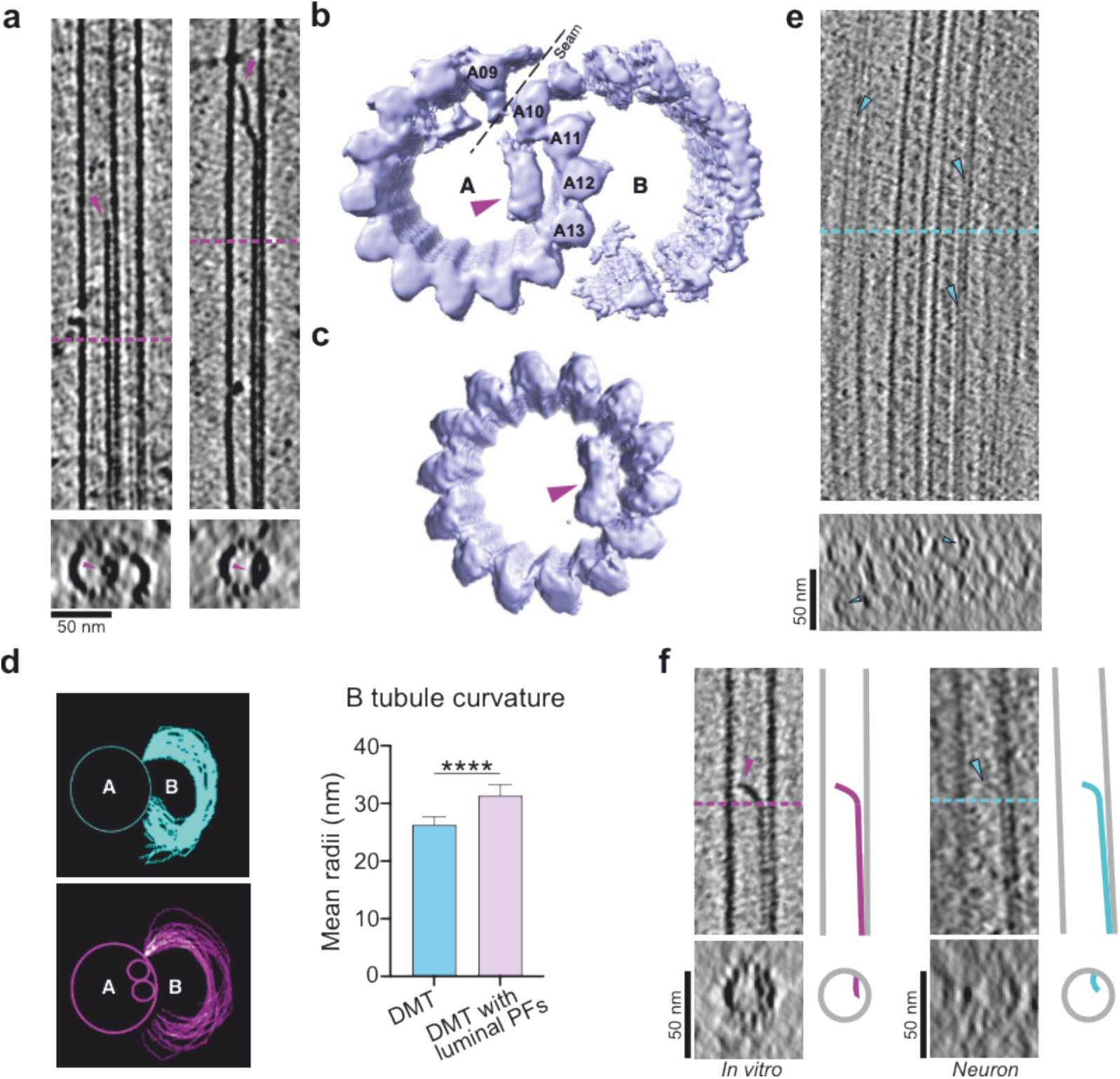
MAP6d1 assembles protofilaments in the microtubule lumen. **a** Examples of tomograms of doublet (left) and singlet (right) microtubules containing luminal protofilaments and their corresponding subtomogram average reconstruction in (**b**) and (**c**), respectively. The putative location of the microtubule seam is indicated by a dashed line. **d** B-tubule curvatures traced for doublet microtubules without (*top*) and with (*bottom*) luminal protofilaments. The graph shows the mean radii of the B-tubule curvatures (n = 36 and 27 for doublet microtubules without and with luminal protofilaments, respectively). Bars represent mean ± SD. ****p<0.0001, Mann-Whitney test. **e** Example of a tomogram showing neuronal microtubules containing luminal protofilaments (blue arrowheads). **f** Comparison of microtubules containing luminal protofilaments assembled with MAP6d1 in vitro (*left*) with those observed in neurons (*right*) and their corresponding schemes. Pink and blue arrowheads show luminal protofilaments in microtubules assembled in vitro and in neurons, respectively.

We next compared the B-tubule curvatures of doublet microtubules with and without luminal protofilaments (**Fig. 5d**) and found that the mean radii of the B-tubules of doublet microtubules with luminal protofilaments increased by ∼15 % (**Fig. 5d**) compared to those without. As luminal protofilaments are localised in the A-tubule right next to where the B-tubule begins, known as the outer junction, they may influence the conformation of the B-tubule.

Because protofilaments within the microtubule lumen have not been previously described, we used cryo-electron tomography to look for the presence of such microtubule architectures in mature neurons (**Fig. 5e** and **Supplementary Fig. 5**). We found singlet microtubule structures containing filamentous densities that resemble the luminal protofilaments induced *in vitro* by MAP6d1 in neuritic extensions (**Fig. 5f**). We propose that these luminal protofilaments could help explain the strong stability of neuronal microtubules.

## Discussion

These data reveal that MAP6d1 plays a specific role in ciliary development by regulating microtubule architecture. Specifically, the brain-specific MAP6d1 induces the assembly of doublet microtubules. This makes MAP6d1 the first microtubule-associated protein demonstrated to independently reconstitute these ciliary structures. We also observed a previously undescribed phenomenon of protofilaments within the lumen that we suspect lend extra strength to neuronal microtubules.

MAP6d1 forms doublet microtubules through tubulin recruitment on the A-tubule lattice, facilitating the nucleation of the B-tubule. MAP6d1 thus behaves both as a MAP, as it can recruit tubulin on the microtubule lattice, as well as a MIP given the presence of its Mn-motifs and its ability to form luminal protofilaments. We propose that this dual binding mode is key to MAP6d1’s unique property to assemble doublet microtubules by bridging the A-tubule to the B-tubule (**Fig. 6**). Given that the C-terminal tail of tubulin plays a crucial role in doublet microtubule assembly^42,43^, it may be that MAP6d1 binding to the A-tubule triggers a conformational change in the tubulin C-terminal tail, promoting tubulin recruitment and subsequent B-tubule nucleation. Although removal of the C-terminal tail of tubulin allows B-tubule nucleation from multiple sites on the A-tubule^42^, MAP6d1 binding is sufficient to restrict B-tubule nucleation to a single A-tubule protofilament. MAP6d1 therefore appears to provide positional information to recruit tubulin at a specific site on the A-tubule lattice. Since deletion of the N-terminal part of MAP6d1 prevents both doublet formation and tubulin recruitment, we suspect the N-terminal participates in this local C-terminal tubulin conformational change.

**Figure 6.**
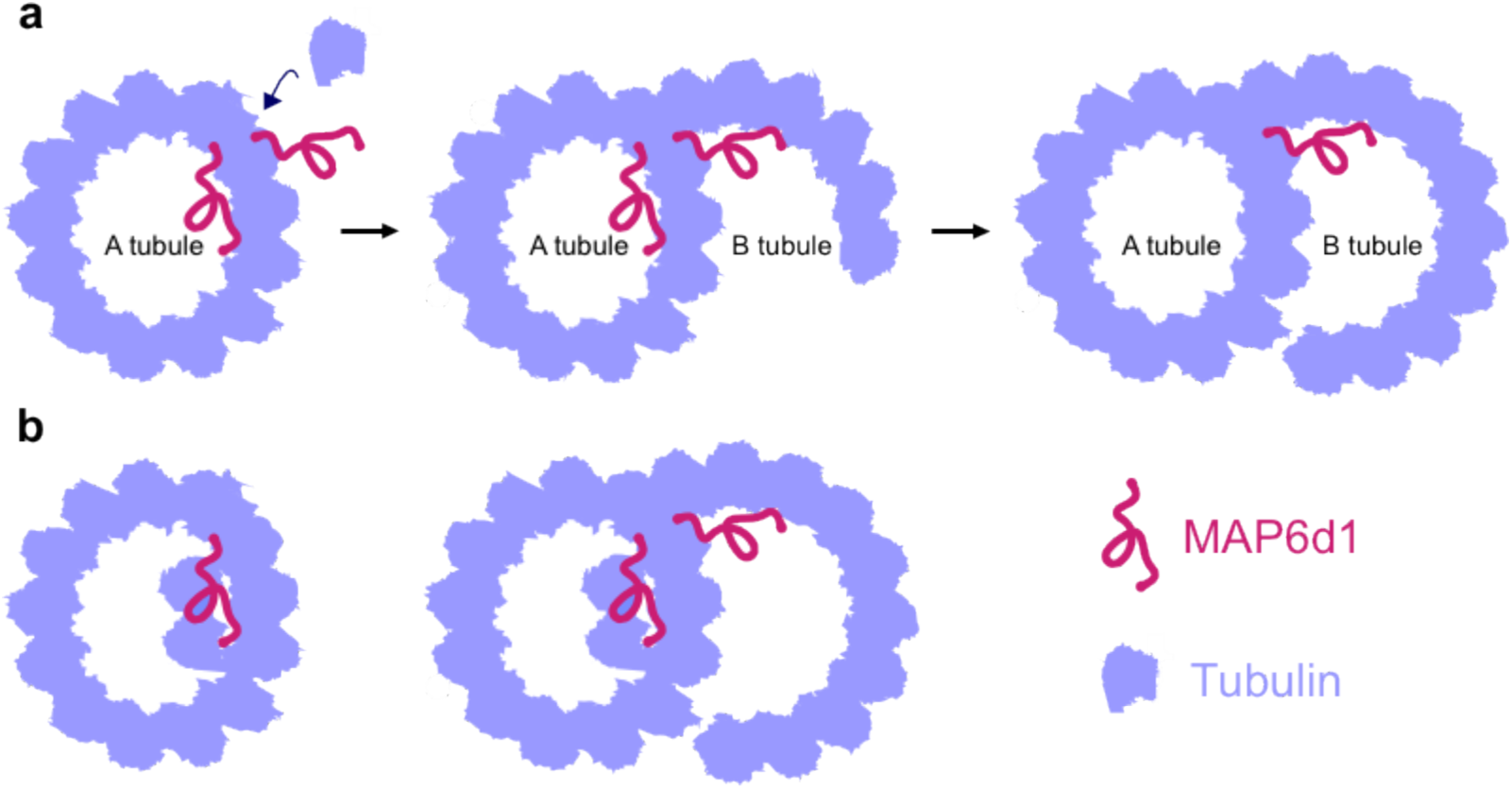
Models of MAP6d1-microtubule interactions leading to the formation of the B-tubule and luminal protofilaments. **a** MAP6d1 binds to the A-tubule lattice, recruits free tubulin dimers to initiate the B-tubule and forms a doublet microtubule. **b** During co-polymerisation with microtubules, MAP6d1 assembles protofilaments in the lumen.

MAP6d1 has one particular feature that may help organise both singlet and doublet microtubules in neurons: it is one of the few proteins, along with the neuronal kinesin-4 (KIF21B) and the ciliary MIP CSPP1, that can induce microtubule pausing^44,45^. It is worth noting in this context that deletion of CSPP1 in zebrafish reduces the length of primary cilia^46^, similar to what we observed in MAP6d1-KO neurons. On the other hand, the MAP6d1-Δ2-35 mutant failed to induce pausing *in vitro* and doubled the length of primary cilia when overexpressed in neurons. It thus seems that microtubule pauses are important for maintaining optimal ciliary length, which varies considerably in response to physiological and pathological changes^47^. Understanding the mechanism by which MAP6d1 induces pauses will require further study, but it could result from a combination of a strong lattice-stabilising effect that prevents tubulin release, as indicated by MAP6d1’s inhibition of microtubule shrinkage, and a modified tip conformation that hinders tubulin incorporation, as proposed for KIF21B^44^.

MAP6d1 may also contribute to microtubule stability in a unique way by assembling protofilaments within the microtubule lumen. In doublet microtubules, luminal protofilaments in the A-tubule always localise near the outer junction, reinforcing the positional specificity of MAP6d1 binding at the B-tubule nucleation site, as discussed above. Such luminal protofilaments could constrain the architecture of the B-tubule or stabilise the doublet outer junction, similar to the organisation of MIPs in axonemal doublets^10,17,48^.

MAP6d1 is part of the SAXO family, which is defined by the universal microtubule lumen-targeting Mn-motif. Most proteins in this family are found within the doublet microtubules of cilia and flagella, where they are thought to stabilise these structures^11,16,17,31,48^. Although structural studies have mapped the distribution of these proteins in doublets, the mechanisms by which they contribute to microtubule stabilisation and/or doublet formation remain largely unexplored. MAP6d1 stands out as the first Mn-motif protein identified in neurons with a specific role in primary cilia and, more generally, as the first protein shown to assemble microtubule doublets. Together, these findings mark an important step toward uncovering the molecular mechanisms driving microtubule doublet formation and understanding the specialised functions of Mn-motif proteins across different cell types.

## Material and Methods

### Plasmid constructs

We cloned cDNAs encoding mouse MAP6d1 and its mutants—MAP6d1-Δ2-35 (deletion of residues 2 to 34), MAP6d1-ΔMn2 (deletion of residues 123 to 145) and MAP6d1-Mn2-7A (amino acids 123 to 128, replaced by alanine), tagged with 8 histidine residues at the C-terminal—in pET42a for bacterial expression. For eukaryotic expression, we used PCR primers for the cloning of mouse MAP6d1-GFP, MAP6d1-Δ2-35-GFP and MAP6d1-ΔMn2-GFP as previously described^30^. Constructs were amplified by PCR on the matrix plasmid MAP6d1 pSG5^30^ with Advantage-GC2 Polymerase Mix (BD Biosciences). The resulting PCR products were cloned into pCR2.1-TOPO (Invitrogen) and then subcloned into pEGFP-N1 (Clontech), in fusion with the EGFP cDNA.

### Tubulin preparation

Tubulin was purified from bovine brain and coupled with biotin, ATTO-565 or ATTO-496 [ATTO-TEC Gmbh, Germany] as previously described^49,50^.

### Recombinant protein expression and purification

MAP6d1 and its mutants were expressed in BL21 (DE3) Star *Escherichia coli* cells. After induction by 0.1 mM IPTG overnight at 18 °C, cells were sonicated in a lysis buffer (50 mM Hepes pH 7.4, 200 mM KCl, 0.1 % Triton, and cOmplete EDTA-free protease inhibitor cocktail tablets [GE, Healthcare]) before centrifugation at 160,000 *g* for 30 min at 4 °C. The clarified lysate was incubated with a cobalt affinity resin (Talon, Clontech) and the protein was eluted by increasing concentrations of imidazole. Collected fractions were concentrated, dialysed against 50 mM Hepes pH 7.4, 200 mM KCl, 20 mM DTT, centrifuged at 100,000 *g* for 5 min at 4 °C before freezing in liquid nitrogen. The concentration was measured using BSA as standard by loading onto an SDS-PAGE gel. Purified proteins used in this study are shown in Supplementary Fig. 1a.

### Preparation of microtubule seeds and Taxol-stabilised microtubules

*GMPCPP-microtubule seeds* (50% biotinylated, 50% ATTO-565 labelled tubulin) were prepared at a final concentration of 10 *μ*M tubulin as previously described^50^. Microtubules were centrifuged at 85,000 *g* for 5 min at 37 °C, resuspended in BRB80 buffer supplemented with 1 mM GMPCPP, aliquoted, and stored in liquid nitrogen.

*Taxol-stabilised microtubules* were polymerised by incubating 70 *μ*M tubulin (50% biotinylated tubulin and 50% ATTO-565 labelled tubulin) in BRB80 buffer (Pipes 80 mM, EGTA 1 mM, MgCl2 1 mM, DTT 1 mM, pH 6.8) with 1 mM GTP for 45 min at 37 °C. 50 *μ*M Taxol was then added and microtubules were incubated for 30 min before being centrifuged at 85,000 *g* for 5 min at 37 °C and resuspended in BRB80 buffer supplemented with 10 *μ*M Taxol.

### Co-sedimentation assay

To assess the microtubule-binding properties of MAP6d1 and its mutants, 1 *μ*M Taxol-stabilised microtubules were incubated for 25 min at 35 °C with 300 nM MAP6d1, MAP6d1-Δ2-35, MAP6d1-ΔMn2 or MAP6d1-Mn2-7A in BRB80 buffer supplemented with 125 mM KCl. The mixtures were then centrifuged at 190,000 *g* (Beckman Coulter rotor TLA-100) for 15 min at 35 °C. Protein contents of the supernatants and pellets were analysed by Western blotting using mouse monoclonal anti-tubulin (clone alpha3A1)^52^ and Penta His HRP-conjugated (Qiagen) antibodies (**Supplementary Fig. 1b**). The intensity of the bands corresponding to MAP6d1 and mutants was quantified using Image J^51^.

### Microtubule dynamics using TIRF microscopy

Chambers made of glass slides and coverslips, functionalised with polyethylene glycol (PEG)-silane and PEG-silane-biotin, respectively, were prepared as previously described^50^. The glass chamber was perfused with neutravidin (25 *μ*g/ml) (Pierce) in 1% BSA in BRB80, followed by Poly(L-lysine) (PLL)-g-PEG (0.1 mg/ml) (JenKem) in 10 mM Hepes, pH 7.4 and flushed with 1% BSA in BRB80. GMPCPP-microtubule seeds were then perfused and incubated for 5 min before being washed twice with 1% BSA in BRB80. Microtubules were polymerised from seeds by adding 12 *μ*M tubulin (15 % ATTO-565 labelled tubulin) with different concentrations of MAP6d1 or its mutants in TIRF buffer (BRB80 containing 1 mM GTP, 25 mM KCl, 3 mM dithiothreitol [DTT], 1 % BSA, glucose [1 mg/ml], catalase [70 *μ*g/ml], glucose oxidase [600 *μ*g/ml] and 0.1 % methylcellulose [4000 cP]). Flow chambers were sealed, and movies were acquired on an inverted microscope (Eclipse TI, Nikon) equipped with an iLas TIRF system (Roper Scientific), a CMOS camera (Prime 95B, Photometrics) and a temperature-controlled stage (LINKAM MC60), under MetaMorph software (version 7.7, Molecular Devices). Samples were excited using a 561-nm laser and observed with an Apochromat 100x oil-immersion objective (N.A., 1.49). Time-lapse acquisitions were carried out for 30 min at 35 °C with a 100 ms exposure and one frame every 2 s.

To assess the effect of dilution of microtubule stability, microtubules were polymerised in glass chambers from ATTO-565-GMPCPP seeds in the presence of 12 *μ*M tubulin (15% ATTO-565 labelled tubulin) and 50 nM MAP6d1 in TIRF buffer containing 1 mM GTP. For control conditions, microtubules were polymerised from 25 *μ*M tubulin in TIRF buffer with 1mM GTP. Unsealed chambers were then incubated for 20 min at 35 °C in a humid environment. Chambers were then placed on the warm stage of the TIRF microscope and microtubule depolymerisation was induced by flowing warm TIRF buffer. Microtubule behaviour was immediately recorded using 561-nm laser and images were acquired every 2 s with a 100 ms exposure for 30 min at 35 °C.

To evaluate the dynamic behaviour of microtubules polymerised at the extremity of MAP6d1-assembled-microtubules, microtubules were grown from ATTO-565-GMPCPP seeds in the presence of 12 *μ*M tubulin (15 % ATTO-565 labelled tubulin) and 50 nM MAP6d1 in TIRF buffer supplemented with 1 mM GTP. The unsealed chambers were incubated for 20 min at 35 °C in a humid environment, before perfusion of a solution containing 12 *μ*M tubulin with 1 mM GTP in TIRF buffer. Images were then acquired under the microscope every 2 s with 100 ms exposure for 30 min at 35 °C.

### Tubulin recruitment assay

Mixtures containing 0.325 *μ*M tubulin (15% ATTO-491 labelled tubulin) and either MAP6d1 (100 nM), MAP6d1-Δ2-35 (100 nM), MAP6d1-ΔMn2 (300 nM) or MAP6d1-Mn2-7A (300 nM) in TIRF buffer with 50 *μ*M GMPCPP and 50 mM KCl were incubated with ATTO-565-GMPCPP seeds in chambers. Samples were excited using 491- and 561-nm lasers, and time-lapse images were acquired on the TIRF microscope every 2 s with 100 ms exposure for 30 min at 35 °C.

### TIRF analysis

Movies were analysed using ImageJ software to generate kymographs of individual microtubules. The four parameters to assess microtubule dynamics were extracted from kymographs using an in-house plugin^51^. Growth and shrinkage rates were calculated from the slopes of the corresponding growth and shrinkage events, respectively. Catastrophe and rescue frequencies were calculated for each microtubule by dividing the number of events by their corresponding growth or shrinkage event durations. Microtubule pauses were defined by growth/shrinkage rates less than 0.24 *μ*m/min at the plus end and 0.12 *μ*m/min at the minus end.

### Negative staining electron microscopy

We used two protocols to evaluate microtubule doublet formation. In one, we co-polymerised 25 *μ*M purified brain tubulin with 250 nM MAP6d1 in BRB80 buffer supplemented with 1 mM GTP and 125 mM KCl for 10 min at 35 °C (**Fig. 2a**). In the second approach, we incubated 10 *μ*M stabilised GMPCPP-microtubule seeds with purified brain tubulin (0.325 *μ*M) and either MAP6d1 (100 nM), MAP6d1-Δ2-35 (100 nM), MAP6d1-ΔMn2 300 nM) or MAP6d1-Mn2-7A (300 nM) in BRB80 buffer supplemented with 50 *μ*M GMPCPP and 50 mM KCl for 10 min at 35 °C (**Fig. 2e**). We then loaded 4 μl of each sample on a 400-mesh copper carbon grid (Electron Microscopy Sciences), washed it briefly with warm BRB80 buffer and stained it with 2.5 % uranyl acetate. Images were acquired using a transmission electron microscope (TEM) JEOL 1200 EX (Veleta). The length of microtubule doublets and singlets was measured on images using ImageJ software.

### Cryo-electron tomography

#### In vitro sample preparation

25 *μ*M tubulin was co-polymerised with 250 nM MAP6d1 supplemented with 1 mM GTP and 125 mM KCl in BRB80 buffer for 5 - 10 min at 35 °C. 4 *μ*l of the solution was mixed to 1 *μ*l of cationic BSA-coated gold beads (Aurion Gold Tracers, 210111) and then loaded on a glow-discharged Quantifoil R2/2 200 mesh copper grid (Electron Microscopy Sciences) and placed in the Leica EM-GP2 95 humidity chamber. The grid was blotted for 2 s and plunge-frozen in liquid ethane.

#### Neuronal sample preparation

glow-discharged Quantifoil R2/2 200 mesh gold grids (Electron Microscopy Sciences) were placed in poly-L-lysine-coated sterile 35-mm glass-bottom dishes (Ibidi, FluoroDish) and hippocampal neurons grown on the above-mentioned grids for 25 days, following the protocol described for neuronal culture. The grid was blotted for 5 s using the vitrification robot Leica EM-GP2 and plunge-frozen in liquid ethane.

#### Image acquisition

The tilt series were acquired using K3 Titan Krios at the ESRF facility^53^ operated at 300 keV, at a magnification of 33,000x between −60° and +60° with a 3° increment under an electron dose of 2.87 e/Å^2^ at a defocus of −3 to −6 *μ*m, using TOMO software (Thermo Fisher Scientific). The acquisition was done with a super-resolution pixel size of 1.35 Å/px and the data processing was carried out in binning 2 with a pixel size of 2.7 Å/px.

#### Image processing

252 and 118 tilt series composed of 41 images were acquired for the in vitro samples and neurons, respectively. Raw tilt images were motion-corrected using MotionCorr2^54^ and then aligned for reconstructing the 3D tomograms by automatic alignment followed by contrast transfer function (CTF) estimation using the tomography pipeline in Eman2 software^55^.

For in vitro samples, we selected microtubules manually using the filament tracing tool and analysed the populations of different microtubule architectures (Supplementary Fig. 3a, b). 4997 particles were extracted from doublet microtubules, with periodicities of 8 nm. Because the doublet microtubule particle population is heterogeneous, with varying numbers of protofilaments and varying curvatures of the B-tubule, we first obtained models with various B-tubules using the particles of individual doublet microtubules (**Fig. 2b**). We then classified the particles according to the number of protofilaments in the B-tubule, using the multi-reference refinement package in Eman2. Particles with 7, 8, 9, 10, or 11 protofilaments in the B-tubule made up 21.27, 11.28, 9.85, 39.92 and 17.67% of the whole population, respectively (**Supplementary Fig. 3c**). Subtomogram averaging refinements were performed on all the extracted particles; we used the doublet microtubule with 10 protofilaments in the B-tubule as the reference, since it was the largest class (39.92%). We generated initial 3D models then applied the new 3D refinement pipeline that includes iterative steps of 3D particle alignment, sub-tilt translational and rotational refinements followed by defocus variation in Eman2. For reconstruction, a mask including both A- and B-tubule was first used to perform subtomogram averaging. We then applied a local mask on the B-tubule to average it independently of the A-tubule. The map displayed in **Figure 2D** is the combination of both maps (**Supplementary Fig. 3d-f**).

From singlet and doublet microtubules with luminal protofilaments, we extracted 5881 and 1680 particles, respectively, with 8 nm periodicity for subtomogram averaging using the new 3D refinement pipeline in Eman2. The number of protofilaments in both singlet and doublet microtubules was analysed on extracted particles by performing 3D classification in Eman2, whereas for the control, this analysis was performed based on the fringe pattern on 2D cryo-EM images^56^ (**Supplementary Fig. 3c**).

The IMOD package was used for displaying tomograms^57^. Electron density maps were visualized with Chimera and ChimeraX^58^. FSC plots were extracted from Eman2 refinement output (**Supplementary Fig. 3**).

### B-tubule curvature analysis

The B-tubules of doublet microtubules were traced on 9 different positions along the z-axis using ImageJ software. The B-tubules were aligned by the A-tubule and the outer junction, then the mean radii were determined along each curve and plotted on the graphs.

### Animals used in this study

All experiments involving animals were approved by the local animal welfare committee (Comité Local CEEA n°44 – APAFIS number 45896-2023111317017897) and in compliance with the European Community Council Directive (directive 2010/63/EU).

The Sl21 (*Map6d1*) cKO mutant mouse line was established at the Institut Clinique de la Souris (ICS) - PHENOMIN (http://www.phenomin.fr). The targeting vector was constructed as follows: A 3 kb 3’ homology arm fragment was amplified by PCR (from 129svpass genomic DNA) and subcloned in an ICS proprietary vector. This ICS vector bore a floxed and flipped Neomycin resistance cassette and a 5’ LoxP site. A 1.9 kb fragment encompassing 628 bps of proximal promoter and *Map6d1* exon 1 (ENSMUSE00000268483) was cloned in a second step. Finally, a 4.2 kb fragment corresponding to the 5’ homology arms was amplified by PCR and subcloned in step 2 plasmid to generate the final targeting construct. The linearised construct was electroporated in 129svpass (ICS derived P1 line) mouse embryonic stem (ES) cells. After G418 selection, targeted clones were identified by 5’ and 3 Long 6range PCRs and further confirmed by Southern blot with a Neo probe (5’ and 3’ digests). Three positive ES clones were injected into C57BL/6J blastocysts. Chimeras were bred with Flp deleter females (in a C57BL/6J genetic background) in order to obtain the cKO allele with no selection cassette. Germline transmission was achieved.

To obtain the plain KO allele, chimeras were bred with Cre deleter females (also in a C57BL/6J genetic background) (**Supplementary Fig. 6**).

Genotyping was performed by PCR on genomic DNA isolated from mouse tails using following oligonucleotides: 5’-GGGGCTTATGCCTGTGGCTATATG-3’ plus 5’- CAGTCCTCCTAGGTGCAGACTG-3’ to detect the mutant allele and 5’- CATGGATTCCAGGGATCCATCTCTC-3’ to detect the wild-type allele.

### Cultured hippocampal neurons

Hippocampal neurons were isolated from the brains of wild-type and *MAP6d1*-knockout (*MAP6d1*-KO) littermate embryos at embryonic day 17.5 (E17.5) and cultured as described by Gory-Fauré *et al*^59^. Briefly, the brains were dissected, and the hippocampi were removed, dissociated, and plated on poly-L-lysine-coated coverslips in 35mm dishes containing DMEM 10% horse serum. After 2 h 30 min, the culture medium was replaced with MACS medium.

In some experiments, neurons were transfected prior to plating using the Lonza Mouse Neuron Nucleofector 2b Kit (program 0-005, following the manufacturer’s instructions). Neurons were electroporated with 4 *μ*g of plasmid encoding control GFP, MAP6d1-GFP, or MAP6d1-GFP deletion mutants immediately following dissociation. To ensure efficient transfection, neurons from one hippocampus were plated per dish.

### Immunofluorescence microscopy and quantification

Cells were fixed in 4% paraformaldehyde, 4% sucrose in phosphate buffered saline (PBS) for 30 min at 37 °C, followed by permeabilisation in 0.2% Triton X-100 in PBS for 2 min at room temperature. Cells were then incubated with primary antibodies for 1h at room temperature. Coverslips were washed 3 times in PBS - 0.1% Tween20 and incubated with conjugated secondary antibodies for 1h in the dark at room temperature. After three final PBS 0.1% Tween20 washes and one with PBS, nuclei were stained using Hoechst (Sigma) in the mounting medium (Dako).

Primary antibodies used included mouse monoclonal anti-Arl13b (1:1000, Addgene), mouse monoclonal anti-polyglutamylated tubulin (1:1000, Coger), rabbit polyclonal anti-acetylated tubulin (1:5000, Millipore), and rat monoclonal anti-tyrosinated tubulin (1:500, clone YL1/2^60^). The secondary antibodies were donkey anti-mouse Cy3, donkey anti-rabbit AF647 and donkey anti-rat (1/500, Jackson Immuno-Research Laboratory).

Images of cilia were acquired on a Zeiss LSM 710 confocal microscope (equipped with a Zeiss AiryScan module), using a 63x oil-immersion NA 1.4 objective and Zen software (Carl Zeiss MicroImaging).

We measured ciliary length using Fiji^51^ on maximal projection of confocal images of cilia, all combined in a single file to facilitate manipulation. Only cells with visible GFP signal at the Golgi and neurites were included (**Supplementary Fig. 4c**). We traced the cilia manually to determine their lengths using the segmented line tool.

For fluorescence intensity measurements, we imported confocal images into Fiji and max-projected the z-planes spanning the cilium. The entire cilium was manually traced from the most proximal end to the distal end with a width of 0.17 *μ*m. We then extracted the intensity profile of the selected region, normalising length and intensity to a 0-100 scale. To compare MAP6d1-GFP localisation to polyglutamylated tubulin, we drew a line along acetylated tubulin ciliary staining and compared the signal of both proteins along this line. Similarly, we drew a line w along Arl13b ciliary staining to compare the respective protein signals of MAP6d1-GFP and tyrosinated tubulin localisation.

## Statistical analysis

All experiments were repeated at least three times. Statistical analysis was performed with Prism GraphPad version 10. Statistical tests and sample sizes are all indicated in the figure legends.

## Data Availibility

All data needed to evaluate the conclusions in the paper are present in the paper and/or the Supplementary Materials. Any additional datasets, analysis details, and material recipes are available from the corresponding authors upon reasonable request.

## Acknowledgements

We thank N. Chaumontel and C. Miscopein for helping protein purification and cell biology experiments. This work used the Photonic Imaging Center of Grenoble Institute Neuroscience (PIC-GIN, University Grenoble Alpes – INSERM U1216) which is part of the ISdV core facility and certified by the IBiSA label. We also used the two EM facilities in Grenoble: the European Synchrotron Radiation Facility (beam time on CM01) and the Grenoble Instruct-ERIC Center (ISBG; UMS 3518 CNRS CEA-UGA-EMBL) with support from the French Infrastructure for Integrated Structural Biology (FRISBI; ANR-10-INSB-05-02) and GRAL, a project of the University Grenoble Alpes graduate school (Ecoles Universitaires de Recherche) CBH-EUR-GS (ANR-17-EURE-0003) within the Grenoble Partnership for Structural Biology. The IBS Electron Microscope facility is supported by the Auvergne Rhône-Alpes Region, the Fonds Feder, the Fondation pour la Recherche Médicale and GIS-IBiSA. We thank the staff members of PIC-GIN and both EM facilities for their assistance. The mice used in this study were bred at the animal facility of IRIG-DRF-CEA (Grenoble, France) which is supported by funding from GRAL, a program of the University Grenoble Alpes graduate school (Ecoles Universitaires de Recherche) CBH-EUR-GS (ANR-17-EURE-0003). We thank the zootechnicians, S. Bama-Toupet and C. Magallon, for their assistance. This work was supported by the Institut National pour la Santé et la Recherche Médicale (INSERM), the Centre National de la Recherche Scientifique (CNRS) and l’Agence Nationale pour la Recherche (ANR-20-CE13-0005-01, ANR-22-CE16-0018-01, ANR-24-CE13-5131-01). D.G. was supported by ANR (20-CE13-0005-01), J.W. by the French MESR Ministry and M. de A. by ANR (ANR-22-CE16-0018-01).

## Author Contribution

Conceptualization: S.G.-F., L.S., I.A. Investigation: D.G., J.W., J.D., C.B., M. de A., G.E., E.D., S.G.-F., L.S. Formal analysis: D.G., J.W., J.D., M. de A., E.D., S.G.-F., L.S., I.A. Resources. G.E., E.D. Writing - Original draft: D.G, L.S., I.A. Writing - review and editing: A.A., D.G, J.W., E.D., G.E., C.B., S.G-F, L.S., I.A. Visualization: D.G., J.W. Supervision: S.G-F, L.S., I.A. Funding acquisition: A.A., I.A.

