## supplemental figures for "The Mn-motif protein MAP6d1 assembles ciliary doublet microtubules"

**Supplementary figures and legends**

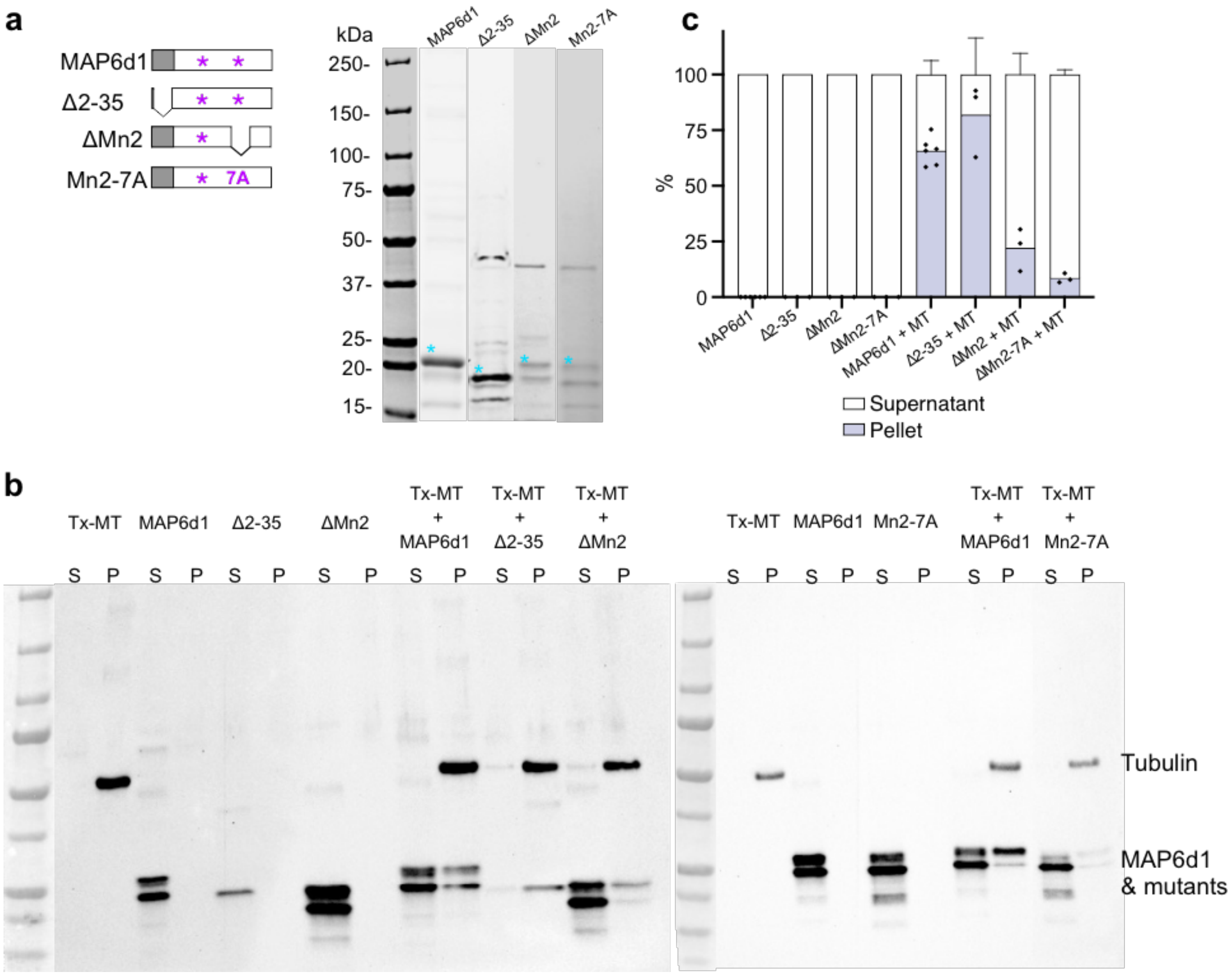


**Supplementary Figure 1. Purified recombinant proteins used in this study and their microtubule-binding properties. a** Purified MAP6d1and the mutants (blue asterisk) on Coomassie-blue-stained gels (10 % SDS PAGE). To the left are diagrams of full-length MAP6d1 and the mutants with the N-terminal in grey and the Mn-motifs with a pink asterisk. **b** Representative immunoblots of co-sedimentation assays of Taxol-stabilised microtubules (Tx-MT) with MAP6d1, and the mutants detected with antibodies against tubulin and histidine (to detect his-tag purified MAP6d1 and mutants). S-Supernatant; P-Pellet. **c** Analysis of blots presented in C. Black dots represent the percentage of proteins in the pellet for each blot. For the two graphs: ****p<0.0001, Fischer's exact contingency test.

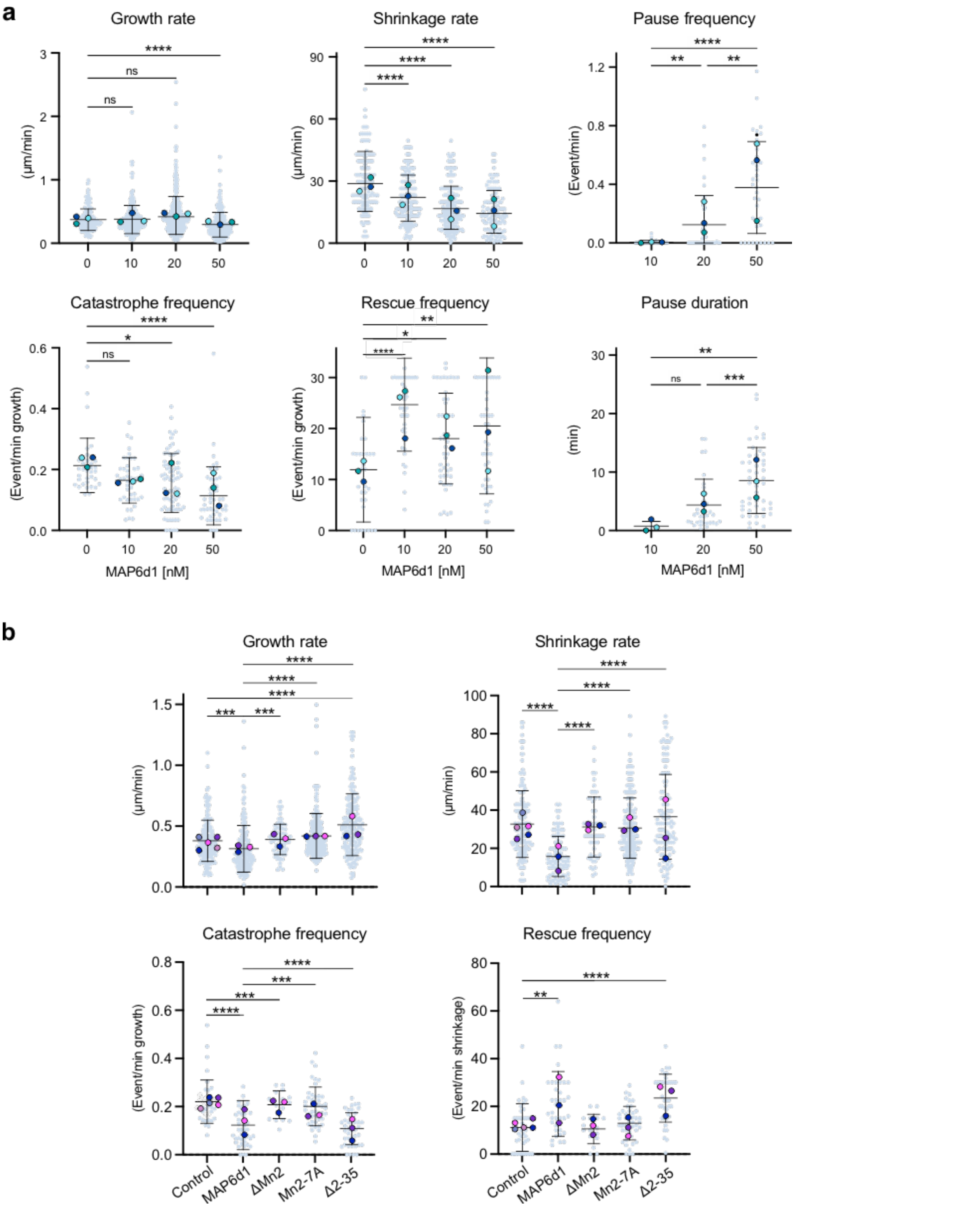


**Supplementary Figure 2. Microtubule minus end dynamic modulation by MAP6d1 and its mutants.**
**a** Graphs showing the growth rate (n = 158, 266, 361, and 217 events for 0, 10, 20 and 50 nM MAP6d1, respectively), shrinkage rate (n = 133, 182, 202 and 116 events), catastrophe and rescue frequencies (n =

35, 43, 72 and 51 MTs), the pause frequencies (n = 31, 41 and 39 MTs, for 10, 20 and 50 nM MAP6d1, respectively) and pause durations (n = 4, 34 and 49 events) of MTs assembled with increasing concentrations of MAP6d1, extracted from kymographs depicted in **Fig. 1a**. **b** Dynamical parameters of microtubules assembled with tubulin alone and in the presence of MAP6d1, MAP6d1-ΔMn2, MAP6d1-Mn2-7A and MAP6d1-Δ2-35: growth rate (n = 187, 175, 68, 239 and 184 events, respectively), shrinkage rate (n = 162, 98, 56, 205 and 117 events, respectively), catastrophe and rescue frequencies (n = 39, 41, 18, 46 and 41 MTs, respectively), extracted from kymographs depicted in **Fig. 3a**. Bars represent mean ± SD from at least three independent experiments. Circles with different colours represent the mean of each individual experiment. *p<0.1, **p<0.01, ***p<0.001, ****p<0.0001, ns = non-significant, Kruskal-Wallis analysis of variance (ANOVA) followed by post hoc Dunn's multiple comparisons tests. For clarity, only significant statistics are indicated for (**b**).

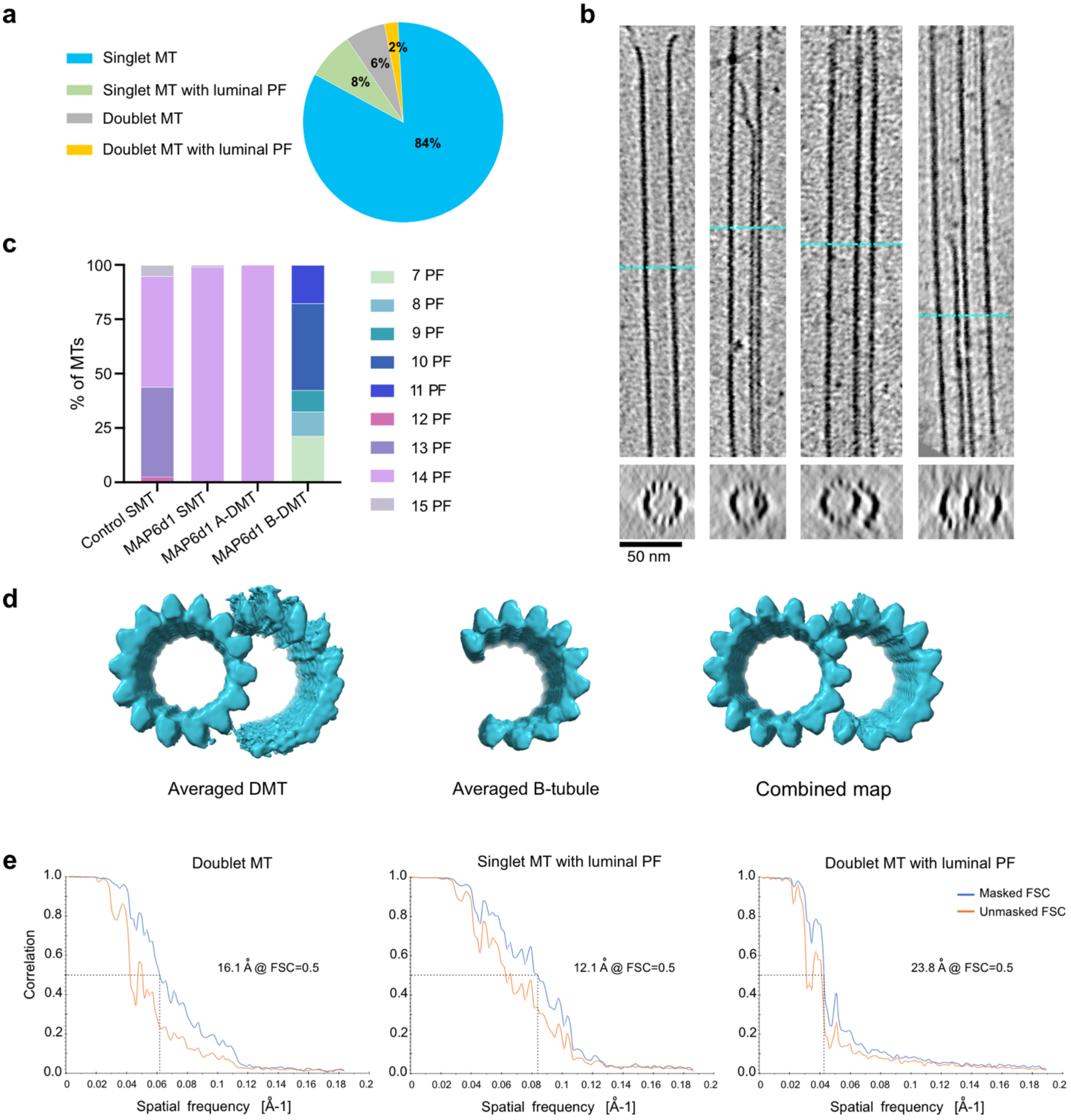


**Supplementary Figure 3. Analysis of MAP6d1-induced microtubule architectures. a** Distribution of microtubule architectures in the tomography data set. **b** Examples of different microtubule architectures induced by MAP6d1. Blue dashed lines indicate the location at which the z-sections are displayed. **c** Distribution of microtubules with different number of protofilaments (PF) measured along 265 $\mu$m of microtubule length for control, and for microtubules polymerised by MAP6d1 are 820 and 115 $\mu$m of SMT and DMT lengths, respectively. **d** Subtomogram averaged models of the initial doublet microtubule, B-tubule and combined map. **e** FSC plots of reconstructions shown in this study obtained from Eman2 refinement output.

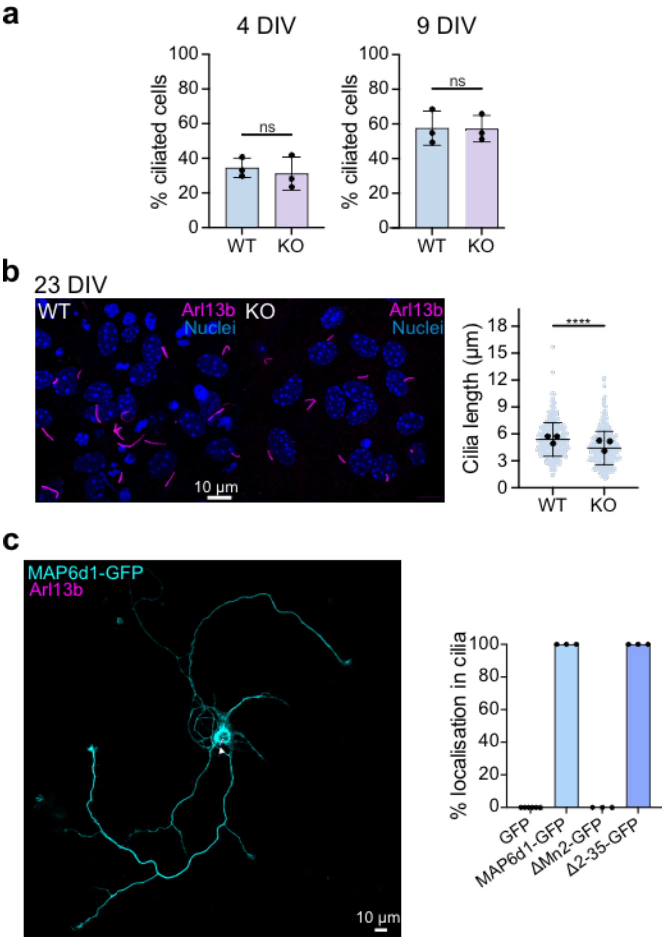


**Supplementary Figure 4. Comparison of primary cilia from wildtype and MAP6d1 deficient neurons. a** Quantification of ciliated cells in hippocampal neurons at 4 days in vitro (DIV) (*left*) and 9 DIV (*right*). n ≥ 350 cells for each individual experiment. The bars show mean ± SD from three independent experiments. Circles with different colours represent the ratio of ciliated cells of each individual experiment. ns = non-significant, Mann-Whitney's test. **b** Representative images of WT and MAP6d1-KO hippocampal neurons at 23 DIV stained against cilia marker Arl13b and with nuclear dye Hoechst. Quantification of cilium length from WT (n = 344) and KO neurons (n = 385), from 3 experiments. Graphs present individual data points with bars representing mean ± SD. **c** Hippocampal neuron at 7 DIV transfected with MAP6d1-GFP and stained against Arl13b. White arrow indicates the primary cilia. The graph represents the proportion of localisation to the cilia in neurons transfected with plasmids encoding GFP (n = 128), MAP6d1-GFP (n = 68), MAP6d1-ΔMn2-GFP (n = 59), MAP6d1-Δ2-35-GFP (n = 86) from at least three independent experiments.

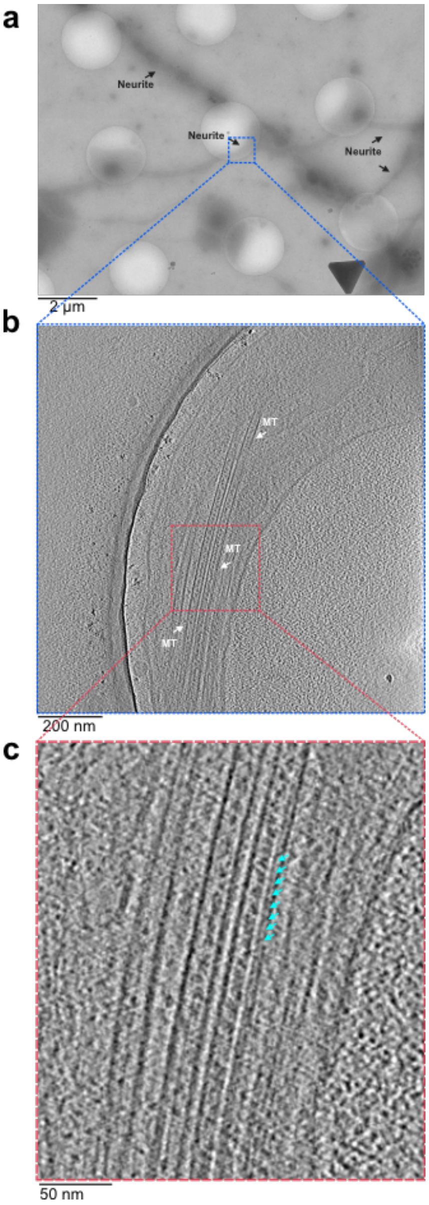


**Supplementary Figure 5. Cryo-electron tomography of microtubules in neurons. a** Low-magnification images of mature neurons on EM grids acquired using Titan Krios cryo-electron microscope. White arrows indicate neurites. **b** High-magnification tomogram of neuritic extensions highlighting microtubules (MTs) with white arrows. **c** Zoomed-in image of the neuronal microtubule with luminal protofilaments indicated by cyan arrows.

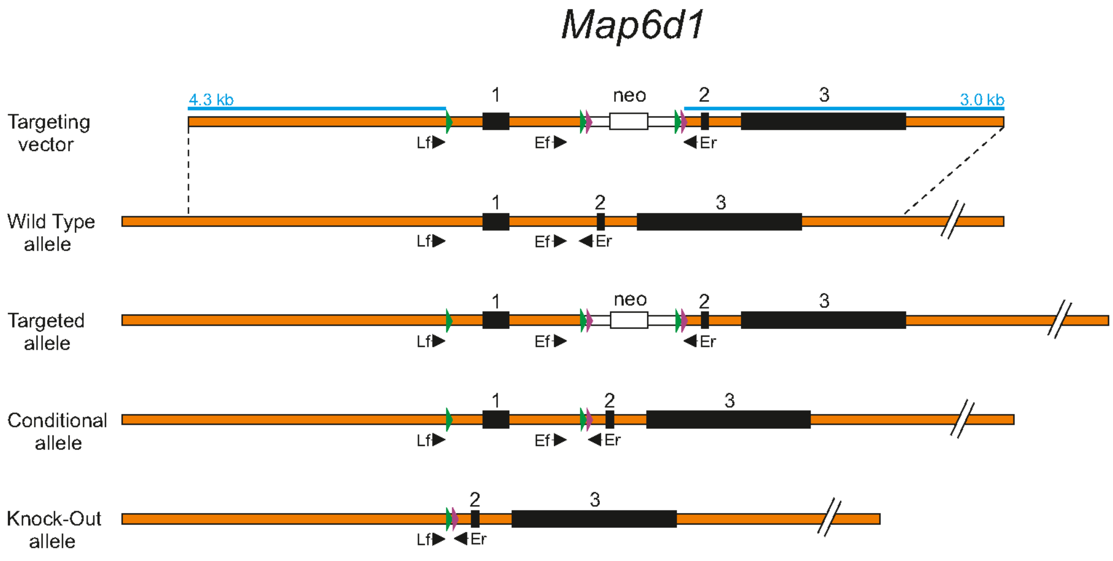


**Supplementary Figure 6. MAP6d1-KO mice generation and genotyping strategy.** Diagram of the targeting vector used and all the possible alleles for *MAP6d1*, and of the wild-type and knock-out alleles for *MAP6d1* (bottom). Orange bar: genomic DNA. Black box: exons with their corresponding number. Green and purple arrowheads: LoxP and Flp sequences, respectively. White bar: neo cassette, with the neomycin resistance gene (white box). Blue lines: zone of sequence homology for homologous recombination, with the corresponding size in kbp. Black arrowheads: primers used for the PCR genotyping.
